# 3D printed cobalt-chromium-molybdenum porous superalloy with superior antiviral activity

**DOI:** 10.1101/2021.07.29.454385

**Authors:** Arun Arjunan, John Robinson, Ahmad Baroutaji, Miguel Martí, Alberto Tuñón-Molina, Ángel Serrano-Aroca

## Abstract

COVID-19 pandemic and associated supply-chain disruptions emphasise the requirement for antimicrobial materials for on-demand manufacturing. Besides aerosol transmission, SARS-CoV-2 is also propagated through contact with virus-contaminated surfaces. As such, the development of effective biofunctional materials that can inactivate SARS-CoV-2 are critical for pandemic preparedness. Such materials will enable the rational development of antiviral devices with prolonged serviceability reducing the environmental burden of disposable alternatives. This research reveals the novel use of Laser Powder Bed Fusion (LPBF) to 3D print porous Cobalt-Chromium-Molybdenum (Co-Cr-Mo) superalloy with potent antiviral activity (100% viral inactivation in 30 mins). The porous material was rationally conceived using a multi-objective surrogate model featuring track thickness (*t*_*t*_) and pore diameter (*ϕ*_*d*_) as responses. The regression analysis found the most significant parameters for Co-Cr-Mo track formation to be the interaction effects of scanning rate (*V*_*s*_) and laser power (*P*_*l*_) in the order *P*_*l*_*V*_*s*_ > *V*_*s*_ > *P*_*l*_. Contrastively, the pore diameter was found to be primarily driven by the hatch spacing (*S*_*h*_). The study is the first to demonstrate the superior antiviral properties of 3D printed Co-Cr-Mo superalloy against an enveloped virus used as biosafe viral model of SARS-CoV-2. The material significantly outperforms the viral inactivation time of other broadly used antiviral metals such as copper and silver from 5 hours to 30 minutes. As such the study goes beyond the current state-of-the-art in antiviral alloys to provide extra-protection to combat the SARS-COV-2 viral spread. The evolving nature of the COVID-19 pandemic brings new and unpredictable challenges where on-demand 3D printing of antiviral materials can achieve rapid solutions while reducing the environmental impact of disposable devices.

**Graphical abstract:** 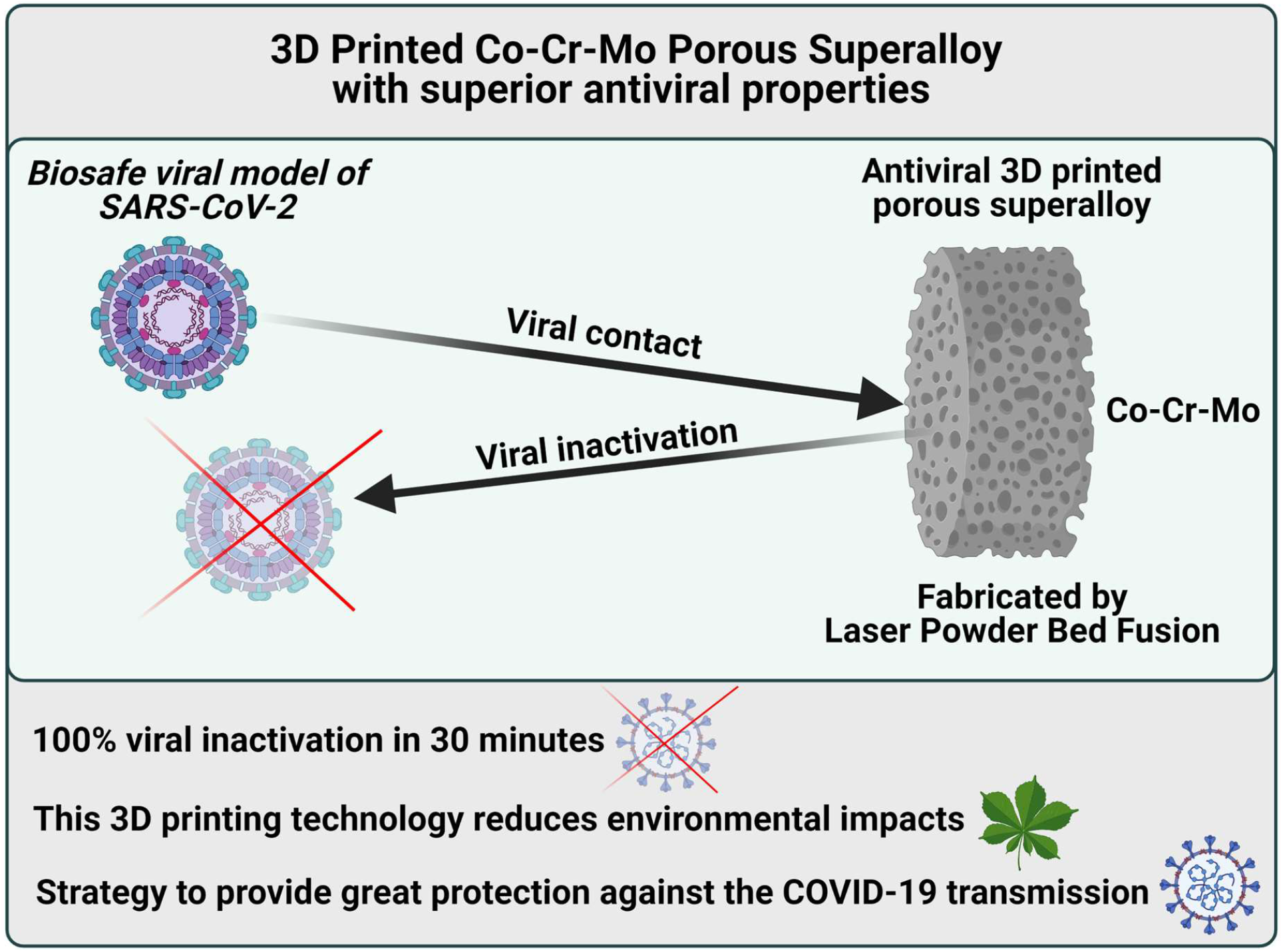

## 1. Introduction

The existence and evolution of SARS-CoV-2 coronavirus variants are spreading more effectively than earlier ones [1,2] due to its evolutionary advantage over SARS-CoV primarily in enhanced host receptor binding [3]. According to Sun and Ostrikov [4], this gives the virus a longer-lasting ability to retain activity on diverse surfaces. As such the lessons so far dealing with SARS-CoV-2 transmission suggests that the new developments of antiviral materials are critical in establishing pandemic preparedness [4,5].

The development of effective materials with antiviral capacity is critical in the development of reusable protective devices such as mask filters, high-efficiency particulate air filters (HEPA) and other antiviral devices [6]. Antiviral portable HEPA filters, air purifiers and aerosol decontaminants are critical also for hospital isolation wards and temporary anterooms [7]. Currently, contaminated devices such as masks and filters are disposable under strict protocol, which is often ignored and possess a serious risk of secondary contamination [8]. A recent study by Maclntyre *et al*. [9] confirmed that even decontamination of reusable masks through washing still offer the probability of infection; further highlighting the need for antiviral materials. Careless disposal of masks is another issue that is resulting in a potential source of microplastic pollution [10] and environmental damage threatening aquatic and animal habitats [11–13].

The development of smart antiviral face mask that contain porous biofilters made of antiviral materials capable of inactivating SARS-CoV-2 is an effective strategy to provide extra protection against the COVID-19 transmission [14] and produce reusable devices to reduce environmental impacts [13,15]. The COVID-19 pandemic has also highlighted the potential for supply chain disruptions causing shortages of essential supplies including PPE, swabs, and ventilators [16– 18]. As such, there is a requirement for on-demand and onsite manufacturing of antiviral materials suitable for a range of applications. Digital manufacturing techniques such as additive manufacturing (3D printing) demonstrated in this study offers significant potential making on-demand and onsite fabrication of antiviral devices accessible [19]. Additive manufacturing (AM) is transforming medical supplies by allowing personalisation and onsite fabrication enhancing resilience against supply chain disruption [20–22]. This subsequently results in rapid development and deployment of potential solutions which is critical when it comes to pandemic preparedness [23–25]. Nevertheless, achieving this requires the development of antimicrobial materials [26–28] and processes that can be additively manufactured without the requirement for complex pre/post-processing as demonstrated in this study.

According to Doremalen *et al*. [29], aerosol and surface transmission of SARS-CoV-2 facilitates infection as the virus can persist for extended periods in a range of common materials as listed in Table 1. Kumar *et al*. [30] highlight that not many studies have examined the effectiveness of metallic materials against SARS-CoV-2 leading to data scarcity for decision making. Generally, the long duration required for inactivation of coronaviruses by metals such as silver (Ag) and copper (Cu) indicates that they might be ineffective when rapid disinfection is required [31,32]. Furthermore, the high cost of these materials is also prohibitive when it comes to their mass adoption as an effective antiviral material.

**Table 1.**
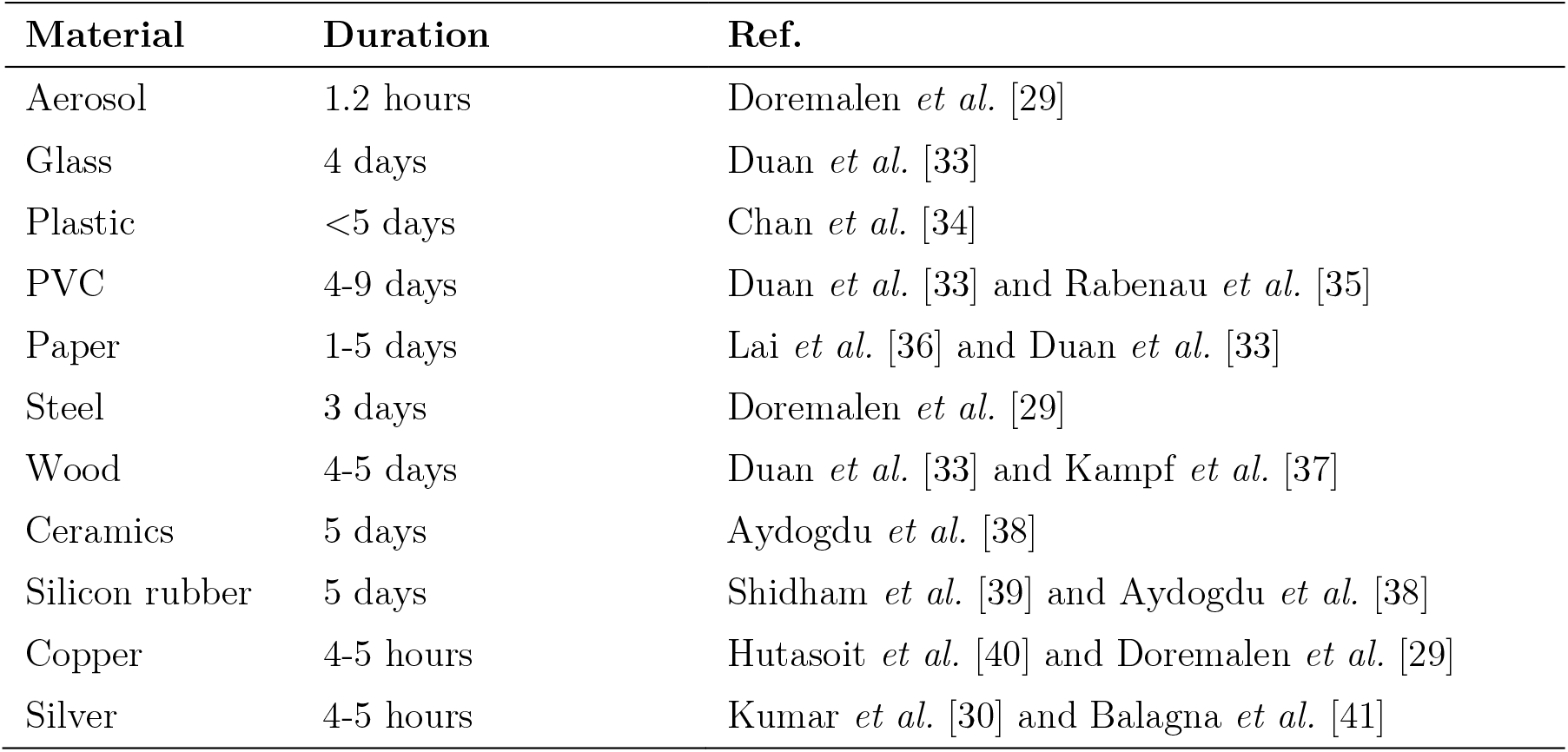
Duration of coronavirus persistence in different materials.

| Material | Duration | Ref. |
| --- | --- | --- |
| Aerosol | 1.2 hours | Doremalen *et al.* [29] |
| Glass | 4 days | Duan *et al.* [33] |
| Plastic | <5 days | Chan *et al.* [34] |
| PVC | 4-9 days | Duan *et al.* [33] and Rabenau *et al.* [35] |
| Paper | 1-5 days | Lai *et al.* [36] and Duan *et al.* [33] |
| Steel | 3 days | Doremalen *et al.* [29] |
| Wood | 4-5 days | Duan *et al.* [33] and Kampf *et al.* [37] |
| Ceramics | 5 days | Aydogdu *et al.* [38] |
| Silicon rubber | 5 days | Shidham *et al.* [39] and Aydogdu *et al.* [38] |
| Copper | 4-5 hours | Hutasoit *et al.* [40] and Doremalen *et al.* [29] |
| Silver | 4-5 hours | Kumar *et al.* [30] and Balagna *et al.* [41] |

When it comes to 3D printed metallic materials, only Cu coating deposited using cold spray have reported 99.2% virus inactivation at 5 hours [40]. Antiviral properties of Cu against SARS-CoV-2 are increasingly being documented at around 4-5 hours [31,42]. Other than Ag and Cu based compositions, studies on the antiviral properties of alternative alloys against SARS-CoV-2 are yet to be reported. As such, the current state-of-art inactivation times for metallic alloys is around 4-5 hours against SARS-CoV-2 which demands composition with superior antiviral efficacy where this study contributes.

According to Hatamie *et al*. [43], cobalt (Co) is an essential trace element that is significantly less toxic in comparison to non-essential metals. Studies on the antimicrobial properties of Co complexes [44] have shown effectiveness against seven microbial strains namely *Bacillus subtilis, Escherichia coli, Klebsiella pneumoniae, Salmonella typhi, Shigella flexneri, Proteus vulgaris* and *Staphylococcus aureus*. Recently, Kota *et al*. [45] showed Co-Cr to be effective in suppressing *S. aureus* and *P. acnes* proliferation both *in vitro* and *in vivo* infection models. Overall, Co-based alloys seem to offer a broad-spectrum antimicrobial possibility making them worthy candidates to explore for potential antiviral activity. Currently, literature on the antiviral properties of Co-based alloys is scarce let alone their performance against SARS-CoV-2.

This research, therefore, is the first step towards drastically improving the state-of-the-art antiviral alloys against SARS-CoV-2 at the interface of 3D printing and surrogate modelling. The research reveals the first Co-Cr-Mo porous material with superior antiviral activity that can be 3D printed on-demand where the innovation pipeline is kept open. The study is directed towards a process-structure–property relationship where both the material and its processing parameters collectively inform an optimum functional architecture. The influence of the LPBF 3D printing process on the structure-property relationship at the sub-micrometre is also analysed to make the digital and on-demand fabrication of the antiviral material accessible and easily scalable.

Overall, the development, analysis, and optimisation Co-Cr-Mo LPBF porous superalloy that feature high antiviral activity against an enveloped virus such as SARS-CoV-2 are demonstrated for the first time. The effect of LPBF process parameters on the characteristics of the porous architecture such as the thickness of the laser melted track (*t*_*t*_) and pore diameter (*ϕ*_*d*_) are also carried out. This was done with the help of a surrogate model that features laser power (*P*_*l*_), hatch spacing (*S*_*h*_) and scanning rate (*V*_*s*_) as LPBF process parameters. The surrogate model was validated and subsequently used for parametric analysis which characterised the order of influence and interaction effects between the process parameters and the resulting printed Co-Cr-Mo architecture.

## 2. Methodology

### 2.1. Laser powder bed fusion (LPBF)

The EOS M290 LPBF machine was used to fabricate the samples at a constant layer thickness of 30 µm. The system featured a 400 W Yb-fibre laser that is modulated above the powder bed within a 250 × 250 mm build platform. Table 2 shows the composition (*wt*. %) of the atomised Co-Cr-Mo alloy used as feedstock at the powder bed. The LPBF process parameters namely the laser power (*P*_*l*_), hatch spacing (*S*_*h*_) and scanning rate (*V*_*s*_) was informed by the surrogate model which is discussed in subsequent sections. The bulk density of the material is 8.3 g/cm^3^ with the morphology of the particles as shown in Fig. 1. The particles featured a spherical morphology with occasional irregular shapes representative of typical feedstock suitable for LPBF. Some smaller satellite particles around 3 µm can be seen attached to larger particles around 18 µm. The overall sphericity and particle size range was found to be favourable for the powder bed fusion process and resulted in even powder spread.

**Table 2.** Composition of the Co-Cr-Mo alloy used as feedstock for LPBF processing.

| Elements | Co | Cr | Mo | Si | Mn | Fe | C | Ni |
| --- | --- | --- | --- | --- | --- | --- | --- | --- |
| Composition (*wt. %*) | 60 – 65 | 26 – 30 | 5 – 7 | $\leq 1.0$ | $\leq 1.0$ | $\leq 0.75$ | $\leq 0.16$ | $\leq 0.1$ |

**Fig. 1.**
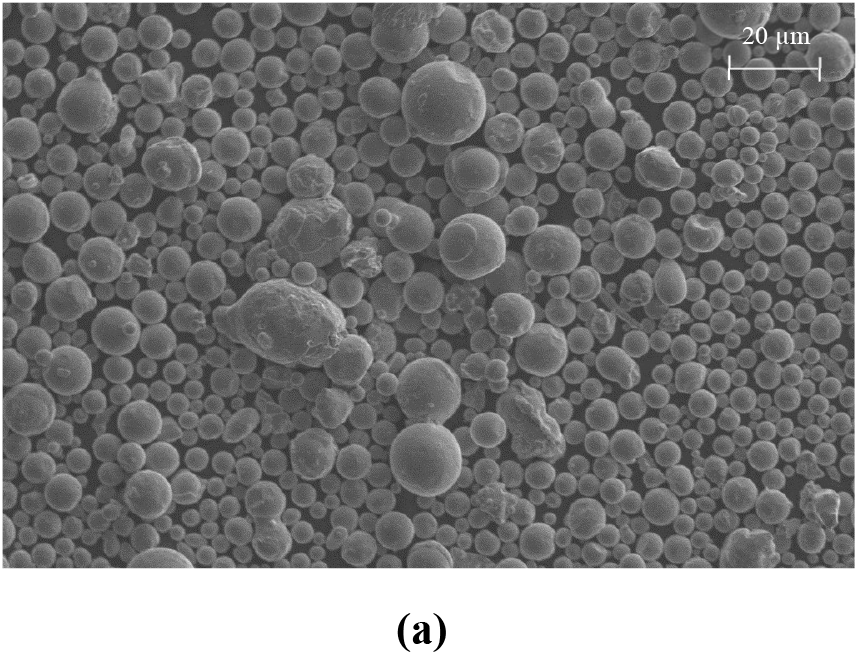
Scanning electron micrograph of atomised Co-Cr-Mo feedstock showing particle morphology.

The laser processing was carried out in an argon inert atmosphere at <0.1% oxygen on a steel substrate of temperature 35° C. The number of samples was informed by the required parametric combinations randomised by the surrogate model. The LPBF process prints follow a layer-by-layer method where the build platform is lowered by the constant layer thickness between subsequent layers. Post-printing the samples were heat-treated at 1050° C for six hours in an argon atmosphere. The resulting track thickness and pore dimensions were characterised using scanning electron microscopy (SEM). EVO 50 SEM produced by Zeiss that uses an incident electron beam to interact with the printed sample to generate backscattered and secondary electrons to create an image of the porous sample is used [48–52].

### 2.2. Surrogate modelling

In LPBF, the extent of material melting and track formation is dictated by the energy density (*e*_*ld*_) [46] at the powder bed which is given by Eq. (1):

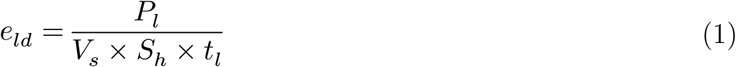

where *P*_*l*_ is the laser power (W), *V*_*s*_ is the scanning rate (mm/s), *S*_*h*_ is the hatch spacing (mm) and *t*_*l*_ is the layer thickness. The relationship between the LPBF process parameters and the energy density at the powder bed means that parameters can be controlled for the targeted outcome. Traditionally, the approach is to identify the optimal energy density required to fully melt and fuse the feedstock resulting in a stable track. Building on this relationship, the study attempts to identify the required LPBF parametric combinations to create Co-Cr-Mo construct with controllable pore size and track thickness. To do this, the relationship between the LPBF process parameters is linked to an objective function of the resulting architecture using Eq. (2):

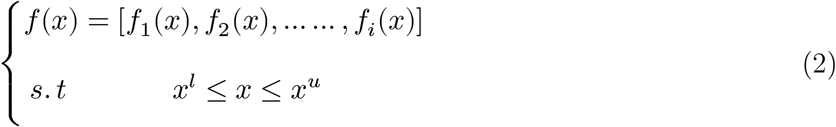

where *x* = (*x*_*l*_, *x*_2_, …, *x*_*k*_) is the vector of *k* LPBF process variables for Co-Cr-Mo. The maximum and minimum limits associated with each process parameters are defined by *x*^*l*^ and *x*^*u*^ for the objective function *f* (*x*). Printing porous Co-Cr-Mo requires characterising the order of influence of the LPBF variables that leads to specific responses. The methodology was to vary the laser energy at the powder bed by algorithmically modifying the LPBF parameters *P*_*l*_, *S*_*h*_ and *V*_*s*_ to inform the track thickness (*t*_*t*_) and pore diameter (*ϕ*_*d*_) as shown in Fig. 2.

**Fig. 2.**
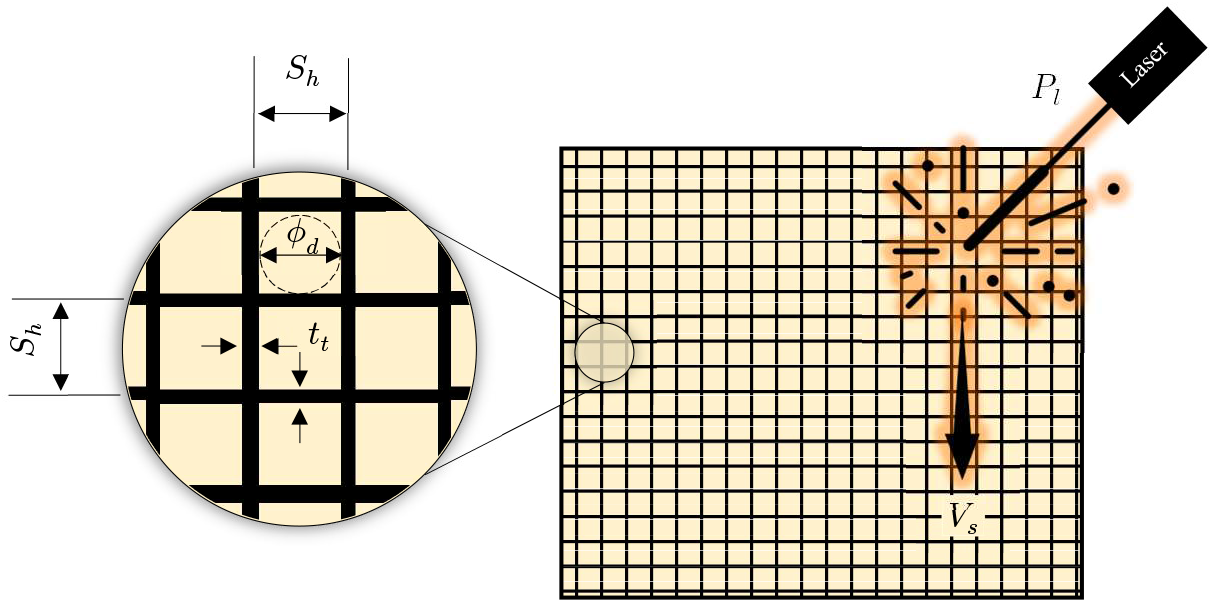
Problem description used for surrogate modelling showing the LPBF variables (*P*_*l*_, *S*_*h*_ and *V*_*s*_) and measured responses (*t*_*t*_ *and ϕ*_*d*_) for the Co-Cr-Mo anti-SARS-CoV-2 porous architecture.

The parametric combination of the randomised training matrix for the surrogate model was conceived based on the Box-Behnken design (BBD) principle. BBD was chosen as it allows for higher-order response surfaces using fewer required runs than a normal factorial technique in the training matrix. As such it is an alternative to the 3^*k*^ factorial that results in an efficient sampling matrix of the coded factorial levels. This approach generally results in a good fit with sufficient information to test for lack of fit. The method also allows models of increasing order to be constructed sequentially and allows for an estimation of experimental error.

The limits of the LPBF process parameters considered for the coded BBD factorial matrix are summarised in Table 3. Based on the problem description, regression analysis was used to characterise the relationship between the LPBF process variables and the resulting responses of the Co-Cr-Mo printed samples. Subsequently, best fit empirical models were derived through randomised experimental data measured for the responses *t*_*t*_ and *ϕ*_*d*_. The resulting polynomial functions were used to predict the order of influence of the contributing process parameters on the responses of the Co-Cr-Mo printed samples.

**Table 3.** LPBF process variables and associated ranges considered for the Co-Cr-Mo surrogate model.

| Variables | Description | Units | Codes | Min. | Med. | Max. |
| --- | --- | --- | --- | --- | --- | --- |
| $S_h$ | Hatch spacing | mm | A | 0.14 | 0.37 | 0.60 |
| $V_s$ | Scanning rate | mm/s | B | 775 | 900 | 1025 |
| $P_l$ | Laser power | W | C | 85 | 110 | 135 |

The steps that led to the development of an accurate surrogate model is summarised in Fig. 3. The surrogate model was trained using randomised test data to satisfy the sampling matrix and analysis of variance criteria. The predictions of the trained model were evaluated for accuracy followed using a desirability criterion to identify the optimum parametric combination that leads to the thinnest Co-Cr-Mo melt tracks and smallest pore diameter. This optimum criterion was chosen to minimise pore diameter and track thickness to accommodate the highest number of pores as possible. The surrogate model was subsequently used to quantify the interaction effects of the process parameters and their influence on the Co-Cr-Mo porous architecture.

**Fig. 3.**
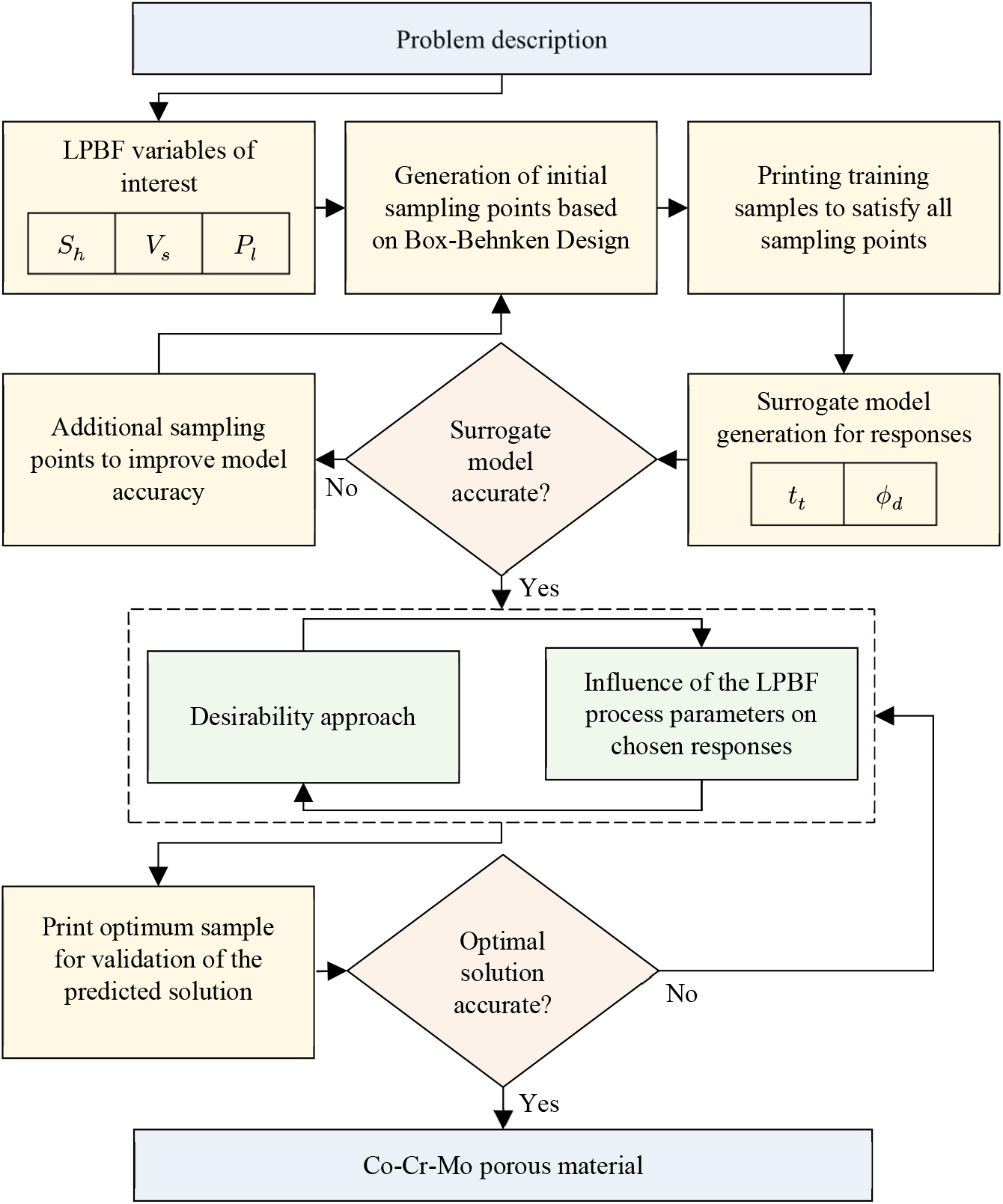
The methodology used to develop the surrogate model for LPBF Co-Cr-Mo porous material.

### 2.3. Antiviral characterisation

The *Phage phi6* host culture was carried using *Pseudomonas syringae* from DSMZ-German collection of microorganisms and cell cultures (DSM 21482) in solid Tryptic Soy Agar (TSA, Liofilchem). Post-culture, the bacteria were incubated in Liquid Tryptic Soy Broth (TSB, Liofilchem) at 120 rpm in 25° C. *Pseudomonas phage phi6* titration was done in accordance with specifications for phage infection.

Phi6 is a double-stranded RNA virus with three-part, segmented, totalling ∼13.5 kb in length. This type of lytic bacteriophage belongs to Group III Baltimore classification and was chosen as SARS-CoV-2 viral model (Group IV: positive-sense single-stranded RNA viruses) as it features a lipid membrane around their nucleocapsid. According to Kitajima *et al*. [47] low pathogenic CoV strains (such as MHV and classical human CoVs) or the same enveloped *Pseudomonas phage phi6* can be used as models of SARS-CoV-2 considering biosafety [48]. Furthermore, recent experiments performed with Phi6 and SARS-CoV-2 have validated the use of this biosafe viral model [14].

LPBF processed Co-Cr-Mo porous samples (*n* = 3) of 10 mm dia. were characterised. A non-woven spunlace fabric filter from NV EVOLUTIA (commercial filters used for face masks) of 10 mm in diameter was cut with a cylindrical punch and used as reference material. All samples were dried at 60° C for 24 hrs. between glass plates and sterilised in ethanol/distilled water (70:30 *vol*. %) solution for 5 mins. at 25° C and UV radiated for an hour on each side.

The antiviral activity of the samples was measured at 30 minutes and five hours of contact with the biosafe viral model. A 50 μL volume of phage suspension in TSB was introduced into each sample at a concentration of 1 × 10^6^ plaque‐forming units per mL (PFU/mL) and incubated for 30 mins and 5 hrs., respectively. Then the samples were placed in a falcon tube with 10 mL TSB and sonicated for 5 mins. at 24 ° C and vortexed for 1 min. Serial dilutions of each falcon were made and 100 μL of each phage dilution were placed in contact with 100 μL of the host strain at OD_600mm_ = 0.5. The infective capacity of the phage was measured by the double-layer method where 4 mL of top agar (TSB + 0.75% bacteriological agar, Scharlau) and 5mM CaCl_2_ were introduced to the phage-bacteria mix. The mixture was poured on TSA plates that were incubated for 24-48h in an oven at 25° C.

## 3. Results and discussion

### 3.1. Morphology LPBF samples

Although Co-Cr-Mo based superalloys are suitable for LPBF [49–51], no studies have demonstrated rationally conceived process informed porosity at track thicknesses below 300 µm. Before identifying the influence of the process parameters on the fabricated samples, the working limits of the LPBF process variables are assessed. This was a critical step as the porosity of Co-Cr-Mo is dictated by the process parameters as opposed to geometry.

The Co-Cr-Mo porous test specimens were printed for all LPBF parametric combinations informed by the rationally conceived surrogate model. The samples were removed from the build plate and analysed under SEM to characterise their porosity. The resulting morphologies of LPBF Co-Cr-Mo porous architecture informed by randomised parametric combinations are shown in Fig. 4. The parametric combinations of laser power (*P*_*l*_), scan speed (*V*_*s*_), and hatch spacing (*S*_*h*_) are sufficient for raising the required energy density at the powder bed for printing Co-Cr-Mo at varying porosity. The entire parametric range selected for the training matrix was found to be suitable to fully laser melt Co-Cr-Mo showing stable and continuous track formation.

**Fig. 4.**
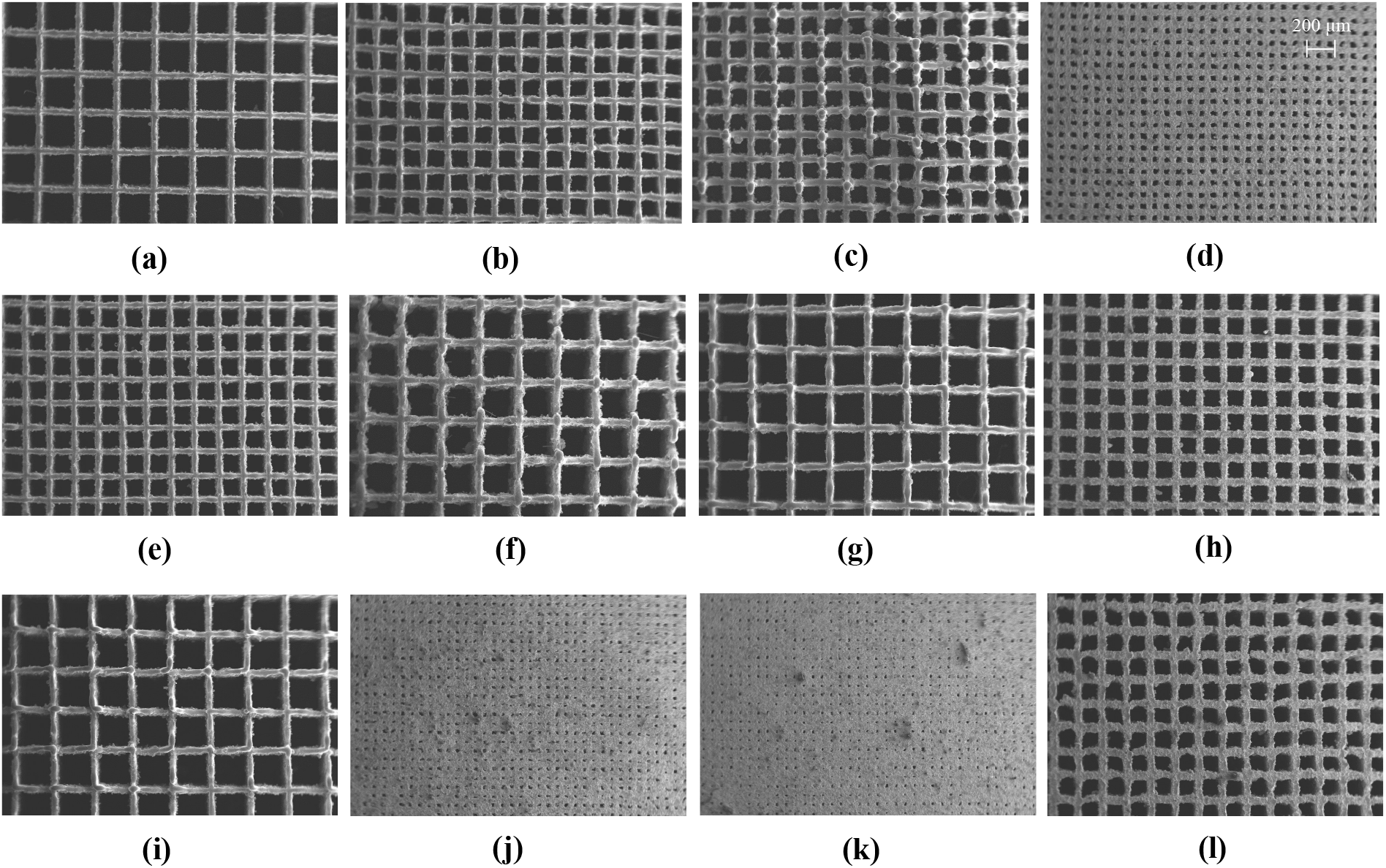
SEM data of additively manufactured Co-Cr-Mo informed by randomised process parameter combinations that were subsequently used to train the surrogate model used to generate optimum Co-Cr-Mo microporous superalloy showing **(a)** *S*_*h*_ = 0.60 *mm, V*_*s*_ = 900 *mm*/*s, P*_*l*_ = 85 *W*, **(b)** *S*_*h*_ = 0.37 *mm, V*_*s*_ = 900 *mm*/*s, P*_*l*_ = 110 *W*, **(c)** *S*_*h*_ = 0.37 *mm, V*_*s*_ = 1025 *mm*/*s, P*_*l*_ = 135 *W*, **(d)** *S*_*h*_ = 0.14 *mm, V*_*s*_ = 900 *mm*/*s, P*_*l*_ = 85 *W*, **(e)** *S*_*h*_ = 0.37 *mm, V*_*s*_ = 1025 *mm*/*s, P*_*l*_ = 85 *W*, **(f)** *S*_*h*_ = 0.60 *mm, V*_*s*_ = 900 *mm*/*s, P*_*l*_ = 135 *W*, **(g)** *S*_*h*_ = 0.37 *mm, V*_*s*_ = 775 *mm*/*s, P*_*l*_ = 85 *W*, **(h)** *S*_*h*_ = 0.60 *mm, V*_*s*_ = 775 *mm*/*s, P*_*l*_ = 110 *W*, **(i)** *S*_*h*_ = 0.60 *mm, V*_*s*_ = 1025 *mm*/*s, P*_*l*_ = 110 *W*, **(j)** *S*_*h*_ = 0.14 *mm, V*_*s*_ = 1025 *mm*/*s, P*_*l*_ = 110 *W*, **(k)** *S*_*h*_ = 0.14 *mm, V*_*s*_ = 775 *mm*/*s, P*_*l*_ = 110 *W*, **(l)** *S*_*h*_ = 0.37 *mm, V*_*s*_ = 775 *mm*/*s, P*_*l*_ = 135 *W*.

As the pore size reduced significantly as shown in Fig. 4j and 4k, some inconsistency in porosity and spatter contamination can be observed. This is due to the high energy density at the centre of the laser spot causing a recoil pressure at the melt pool while other parts of the melt pool are solidifying, which result in the expulsion of the molten material. The spattered metal subsequently cool down forming particles of varying sizes depending on the duration of the condensation process. These spatter formations are not unique to Co-Cr-Mo and are widely observed in a range of materials processed using powder bed fusion as summarised by Young *et al*. [52]. Although some spatters are observed, these are not extensive and can be seen to not obstruct the overall porous architecture being generated.

Although the printed samples established the suitability of conceiving process-induced porosity, the ideal combination of the process parameters that will result in an optimum porous construct to maximise surface contact is required. However, the parameters should ensure that the generated *ϕ*_*d*_ is not so low making the powder removal impossible. To establish such an optimum parametric combination, the order of influence of the process parameters on the resulting track thickness and pore size is required. This gives rise to a multi-objective optimisation problem which was established using a surrogate model.

### 3.2. Surrogate model

#### 3.2.1. Training matrix and regression analysis

Controlling the three parameters for the formation of targeted track width, and pore size requires establishing a process-property relationship. The attempt is to fabricate Co-Cr-Mo architecture that features the thinnest track width and smallest possible distance between adjacent tracks while preserving porosity. Achieving this requires characterising both the interaction effects and order of influence of the LPBF process variables on the printed Co-Cr-Mo porous material.

To achieve the process-property relationship, a randomised BBD training matrix is generated as shown in Table 4 informed by the factors identified in Table 3. Keeping the layer thickness constant, three remaining primary LPBF process parameters that can vary the energy density at the powder bed were used as the variable factors. Consideration was also given when selecting the maximum and minimum limits of these process parameters to make sure sufficient energy density was available for Co-Cr-Mo melt pool generation.

**Table 4.** Co-Cr-Mo surrogate model training matrix showing randomised parameters and the measured responses.

| Variable factors | | | Responses ($\mu\text{m}$) | |
| --- | --- | --- | --- | --- |
| $A = S_h$ (mm) | $B = V_s$ (mm/s) | $C = P_l$ (W) | $t_t$ | $\phi_d$ |
| 0.60 | 900 | 85 | 97 | 502 |
| 0.37 | 900 | 110 | 118 | 262 |
| 0.37 | 1025 | 135 | 111 | 272 |
| 0.37 | 900 | 110 | 118 | 262 |
| 0.37 | 900 | 110 | 118 | 262 |
| 0.14 | 900 | 85 | 97 | 62 |
| 0.37 | 1025 | 85 | 94 | 264 |
| 0.60 | 900 | 135 | 128 | 479 |
| 0.37 | 775 | 85 | 99 | 275 |
| 0.60 | 775 | 110 | 113 | 474 |
| 0.60 | 1025 | 110 | 78 | 494 |
| 0.37 | 900 | 110 | 118 | 262 |
| 0.37 | 900 | 110 | 118 | 262 |
| 0.14 | 1025 | 110 | 78 | 65 |
| 0.14 | 775 | 110 | 113 | 53 |
| 0.37 | 775 | 135 | 136 | 251 |

Test prints were carried out in Co-Cr-Mo for all parametric combinations dictated by the matrix. Their characteristic results measured using SEM for track thickness and pore diameter are listed in Table 4. The results of the analysis were used to identify LPBF process parameters that had the most and least significance on the responses of interest. While the ideal combination of LPBF process parameters for fully dense Co-Cr-Mo is being increasingly documented [53–55], the literature is rather scarce when it comes to processing porosity and the fabrication of tracks with thickness below 300 µm.

The regression analysis of the training data from Table 4 based on best-fit indicators revealed that the thickness of Co-Cr-Mo track has a quadratic relationship with the LPBF process parameters as listed in Eq. (3). The pore size or the linear distance between two adjacent tracks however was found to follow a linear relationship as listed in Eq. (4). A quadratic dependency when it comes to track thickness indicates strong interaction effects between the LPBF process parameters considered. This means that specifying each of the LPBF process parameters require a critical understanding of the linking parameters and their interaction effects to accurately control the resulting *t*_*t*_.

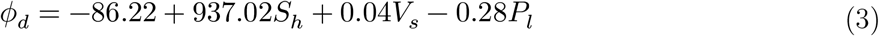

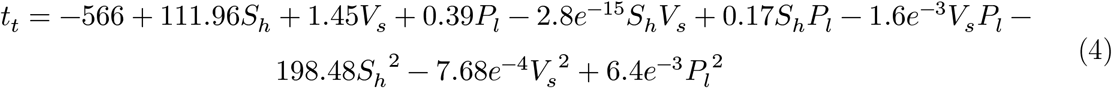

#### 3.2.2. Model accuracy

The accuracy of the surrogate models was evaluated using the analysis of variance (ANOVA) technique where the significant model terms are as summarised in Table 5. The relevant accuracy indicators include the probability (p-value), coefficient of determination ***R***^2^, Adjusted ***R***^2^, and Adequate precision. It can be seen that all models exhibit high F-values and very low p-values confirming that the models are significant. In statistical terms, surrogate models with a p-value less than 0.05 and an adequate precision ratio greater than 4 signifies an accurate model [56]. A closer to unity ***R***^2^ and *Adj-R*^*2*^ also indicates that the surrogate model is accurate for all the responses analysed.

**Table 5.** Analysis of variance showing the accuracy of the surrogate model developed.

| Model | F-value | p-value | Statistical measurements |  |  |
| --- | --- | --- | --- | --- | --- |
| | | | $R^2$ | $Adj-R^2$ | $Adeq-precision$ |
| $t_t$ | 12.60 | 0.003 | 0.9498 | 0.8744 | 12.7989 |
| $\phi_d$ | 1602.67 | <0.0001 | 0.9975 | 0.9969 | 108.7482 |

The relationship between actual responses and those based on the surrogate model are shown in Fig. 5. As can be seen, the residuals are close to the predicted results which validate the accuracy of the surrogate model. Overall, the ANOVA demonstrates that all the models developed in this study are suitable for making valid predictions. This means that Eq. (3-4) adequately characterise the relationship between the laser power, scanning rate, and hatch spacing and to that of the resulting properties of porous Co-Cr-Mo. This means that the surrogate model can be used to analyse the interaction effects between the process parameters and to identify their optimum combinations to print an optimum porous architecture.

**Fig. 5.**
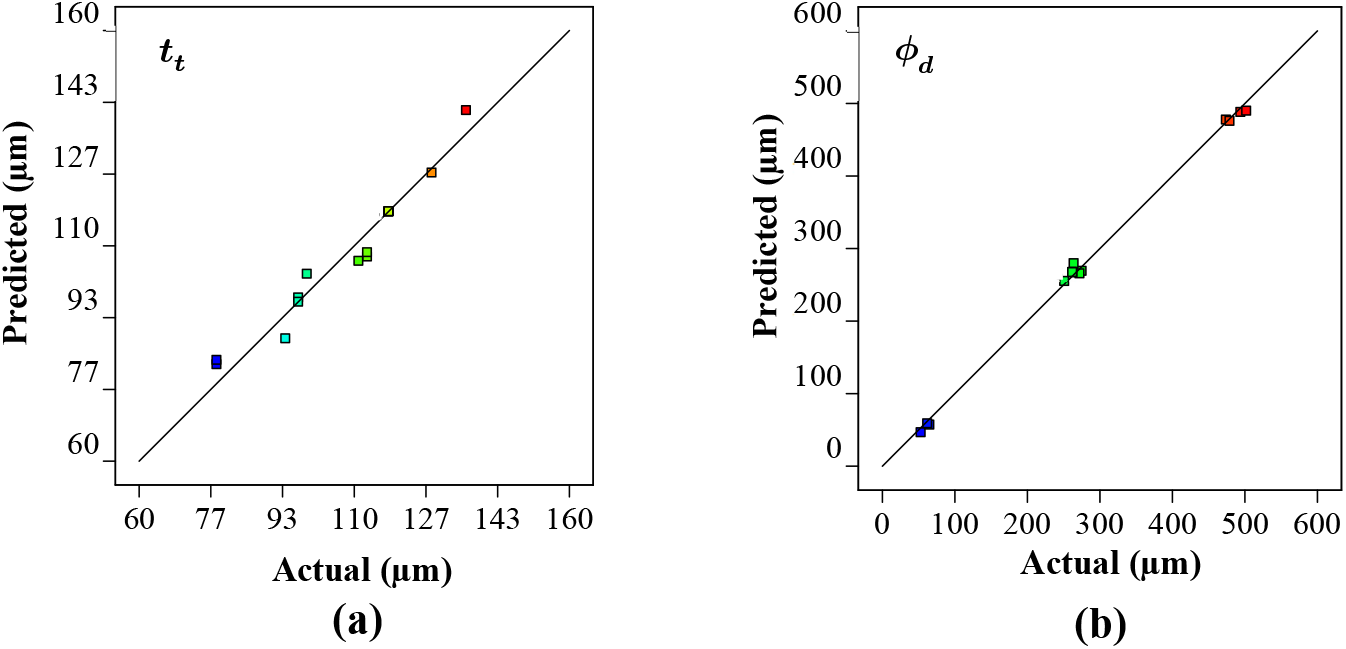
Comparison of experimentally measured and predicted values for the parametrically developed Co-Cr-Mo architecture demonstrating the accuracy of the surrogate model for (a) track thickness, (b) pore diameter.

### 3.3. Significance of individual process parameters

#### 3.3.1. Laser power

When the laser scans over the powder layer, the absorbed energy from the laser beam heat the Co-Cr-Mo particles creating a melt pool. Heating, time-evolution of the melt pool, and the solidification process depend on the powder and process characteristics [57]. The process parameters influence the phases, recoil pressure, surface tension, Marangoni effect, and hydrodynamics that affects the size, and shape of the melt pool. When the laser beam leaves the melt area, the melt pool starts to cool down and solidifies [58,59]. As such the creation of a stable melt pool with a regular shape and geometrical characteristics, the powder characteristics of Co-Cr-Mo and laser processing parameters are the significant factors. These parameters also have a direct influence on the manufacturing time and quality of the final parts.

Fig. 6 shows the Co-Cr-Mo track thickness and pore diameter at varying laser power at a constant hatch spacing and scanning rate. As the laser power increases, the amount of interacting powder rises increasing the track thickness as shown in Fig. 6a. At higher laser power, the Co-Cr-Mo track formation is contributed both from the particles directly and adjacent to the laser spot resulting in a denudation zone. The powder denudation zone defines the volume of powder involved in the track formation and spattering process. The process is not unique to Co-Cr-Mo and is similar to most metals when processed using LPBF [60].

**Fig. 6.**
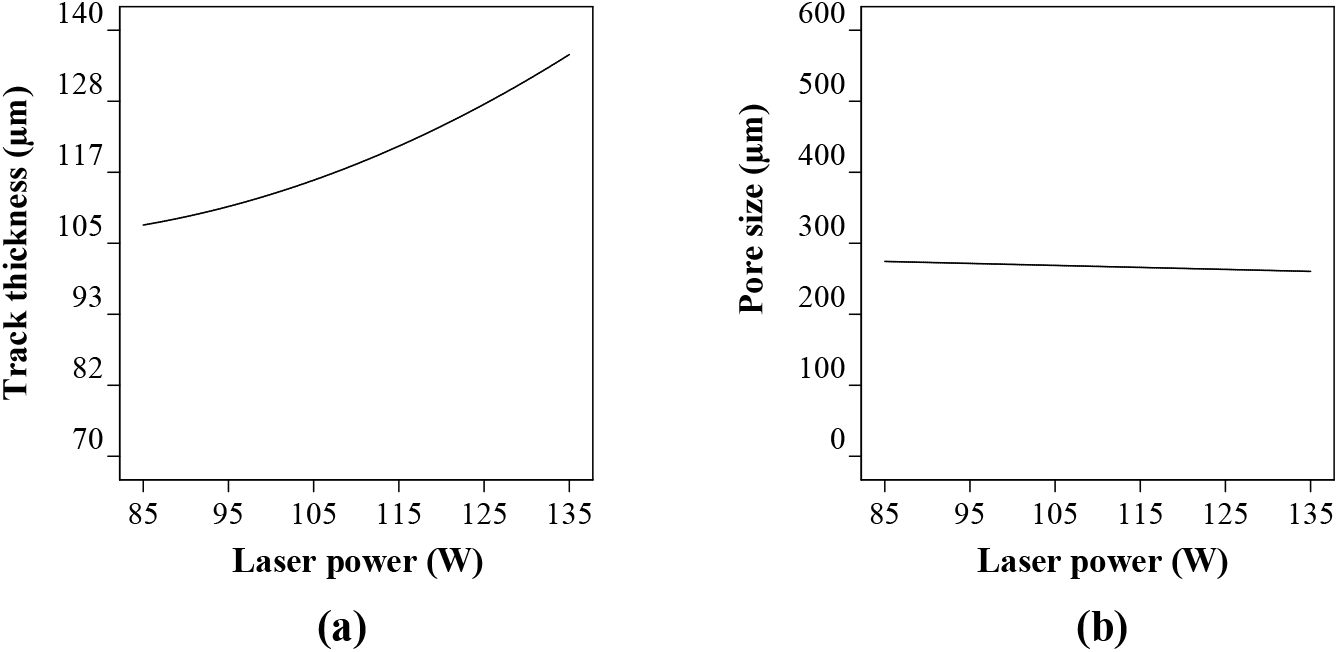
Influence of laser power on (a) the track thickness and (b) pore size of the porous Co-Cr-Mo.

Although there is a consistent increase in ***t***_***t***_ with laser power, the rate of change was found to reduce gradually post 115 W. This is because the rise in laser energy, melt convection, and thermal diffusion increases the unusable energy which cannot be completely absorbed in the powder bed. Consequently, the track thickness and laser power can be observed to have a significant influence but without an absolute proportionality throughout the laser power range considered; which is consistent with the literature [55,61,62]. Overall, a variation in laser power from 85 W to 135 W was found to increase the track thickness by ∼26%.

When it comes to the influence of laser power on the pore size (Fig. 6b), a linear relationship was observed. It was found that the pore size was highest at 274 µm when the laser power was at its lowest signifying a thinner track. As the laser power increased to 135 W, the pore diameter reduced by 5% to 260 µm. As such, without considering any interaction between the process parameters, laser power has the highest influence on the track thickness, which subsequently affects the pore diameter to a lesser extent.

#### 3.3.2. Scanning rate

The scanning rate is the speed at which the laser spot travels across the powder bed. The first consideration when choosing the parameter is to achieve a consistent, fully dense track. For any given build, the powder characteristics and layer thickness are constant parameters. As such using a fixed laser spot size leaves laser power, scanning rate and hatch spacing as the process variables. Generally, all three of these parameters required to be carefully controlled; for example, a high scanning rate can result in insufficient energy at the powder bed. This will lead to an un-melted or partially melted track featuring ‘lack of fusion’ porosity. In contrast, lowering the scanning rate leads to a high energy density that can overheat the melt pool, causing deeper energy penetration leading to keyhole formation and unstable track. Therefore, there is an optimum scanning rate window that achieves continuously fused material track. A fully dense material track is generally the aim while laser processing, however for this study the target is to identify the thinnest most stable track.

The effect of scanning rate on the track thickness and the pore shape were analysed at a constant laser power and hatch spacing of 110 W and 0.37 mm respectively. The entire scanning rate range resulted in fully melted tracks. However, at a low scan rate, the laser interacts with the powder longer resulting in a thicker track. It can be seen from Fig. 7a that the thickness of the molten Co-Cr-Mo track decreased as the scanning speed increased. However, the track thickness shows that the scanning rate has a threshold character characterised by the flattening of curves at a scanning rate around 775-875 mm/s (Fig. 7a).

**Fig. 7.**
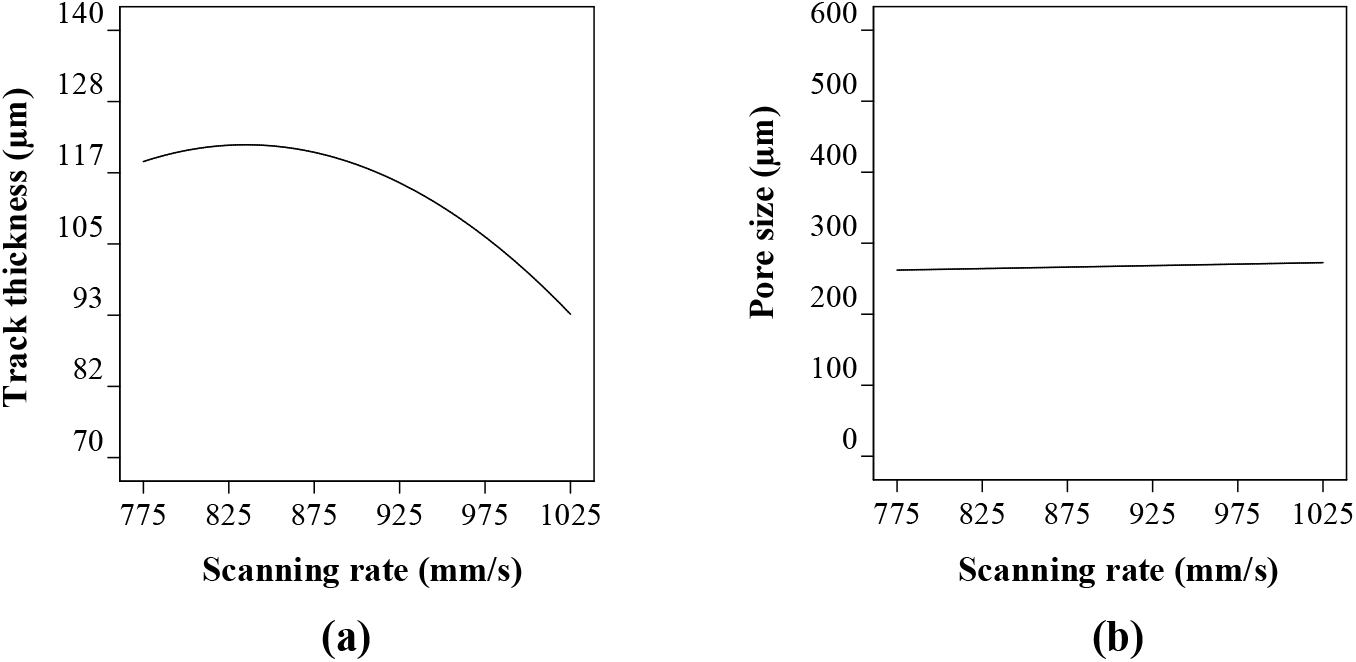
Influence of scanning rate on (a) the track thickness and (b) pore size of LPBF processed Co-Cr-Mo.

Studies by Bayat *et al*. [63], King *et al*. [64] and Robinson *et al*. [65] shows that excessively low scanning rate can result in keyhole mode of melting where the interaction of surface tension and hydrostatic force versus vapour pressure is not balanced, keyholes appear that affect the surface quality. This can be attributed to the increase in temperature of the melt pool at a low scanning rate resulting in stronger Marangoni’s convection and fluid flow creating keyhole pores intermittently. This result in stronger movement of un-melted powders toward the melt pool, resulting in increased roughness and track thickness.

Overall, the highest track thickness of 121 µm was observed at a scanning rate of 825 mm/s which reduced by 22% when the scanning rate was increased to 1025 mm/s. This is because as the scanning rate increases, the residence time of the laser per unit area reduces engaging in a lesser number of powdered particles leading to smaller melt-pool width. Nevertheless, Fig. 7b shows that the changing track thickness has a small influence on the pore size where the diameter of the pore increased by ∼4% when the scanning rate was increased from 775 mm/s to 1025 mm/s. Similar to the case of laser power, the scanning rate was found to have a significant effect on the track thickness which subsequently affected the pore size to a lesser extent. As such, both the laser power and scanning rate did not directly affect the spacing between the melted tracks leading to a lower influence on pore size.

#### 3.3.3. Hatch spacing

The next level of complexity is the melting of multiple tracks to fill the designated part area based on the design. The part area is build-up of adjacent tracks, where their overlap distance is dictated by the hatch spacing. It is measured from the centre of one laser spot to the adjacent.

To have process-induced porosity such as the ones targeted in this study, a large hatch spacing is required. Otherwise, the laser tracks will overlap resulting in a fully dense part.

Fig. 8 shows the influence of hatch spacing when the laser power and scanning rate are kept constant. It can be seen from Fig. 8a that lower and higher hatch spacing seem to have no significant effect on the track thickness. This was expected as the hatch spacing range was deliberately chosen to not coincide or overlap with the previous track to induce a porosity. Nevertheless, a small rise in track width can be observed around a 0.37 mm hatch spacing which can be attributed to the effect of the powder particle size and some inconsistencies in the track formation. Only the analysis on the interaction effect will reveal the exact reason which is discussed in subsequent sections. From Fig. 8b, it is evident that even at the smallest hatch spacing of 0.14 mm there exist a small gap between two consecutive laser tracks. This explains why the track thickness is unaffected despite the change in hatch spacing in Fig. 8a.

**Fig. 8.**
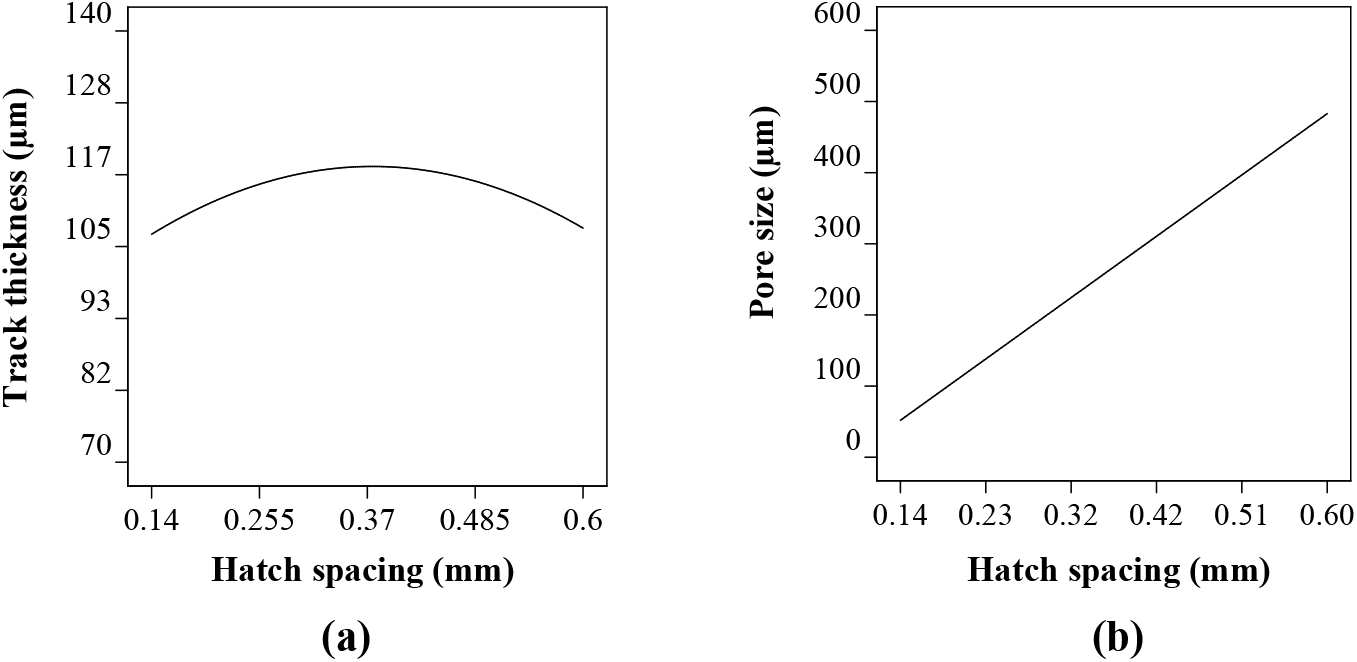
Influence of hatch spacing on (a) the track thickness and (b) pore size of LPBF processed Co-Cr-Mo.

Evaluating Fig. 8b, the hatch spacing has a significant effect on the pore size. The increase in pore diameter is linearly consistent with an increase in hatch spacing. The lowest hatch spacing was found to be 50 µm at a hatch spacing of 0.14 mm, however, this increased almost 10 folds when the hatch spacing was increased to 0.60. Therefore, when it comes to pore size, hatch spacing has the most significant effect despite keeping all other parameters constant. The track thickness on the other hand was primarily affected by the laser power and scanning rate with almost no influence from the hatch spacing range considered. Although the study so far has identified the influence of individual process parameters on the characteristics of the Co-Cr-Mo microporous material, the interaction effects are not understood, which is discussed in subsequent sections.

### 3.4. Interaction effects between LPBF process parameters

#### 3.4.1. Track thickness

Laser power, scanning rate, and hatch spacing are the three variables considered in this study. Although these parameters can be varied independently, their interaction has the most significant effect on the melt pool characteristics. Therefore, studying the interaction effects between the process parameters and identifying their order of influence is critical in identifying the optimum parametric combination. The thickness of Co-Cr-Mo track has a significant influence on the overall pore size and the surface area achievable. Ideally, a stable but thinner track offers a higher opportunity for functional porosity as opposed to thicker tracks. As such identifying and accurately characterising the interaction effects between the LPBF process parameters is carried out to inform the optimised microporous material.

The interaction effects between the process parameters affecting the Co-Cr-Mo track thickness are shown in Fig. 9. As can be seen, the parameters behave differently depending upon their combination with another parameter being used. Looking at the influence of scanning rate and hatch spacing as shown in Fig. 9a; track thickness is slightly decreased as the scanning rate increases resulting in the thinnest track at the highest scan speed with minimal influence from hatch spacing. As such the interaction effects between scanning rate and hatch spacing on Co-Cr-Mo track thickness is insignificant, which is consistent with the single parameter observations. This was expected as the hatch spacing range was deliberately chosen for porosity rather than to allow melt pool overlap. A similar trend was observed regarding the interaction effects between laser power and hatch spacing as shown in Fig. 9b. Here, the thickness of the track is primarily driven by the laser energy with negligible interaction from hatch spacing. As such ***t***_***t***_ increases consistently with the intensity of the laser power. Overall, both Fig. 9a and 9b show that the hatch spacing has minimal interaction when it comes to track thickness.

**Fig. 9.**
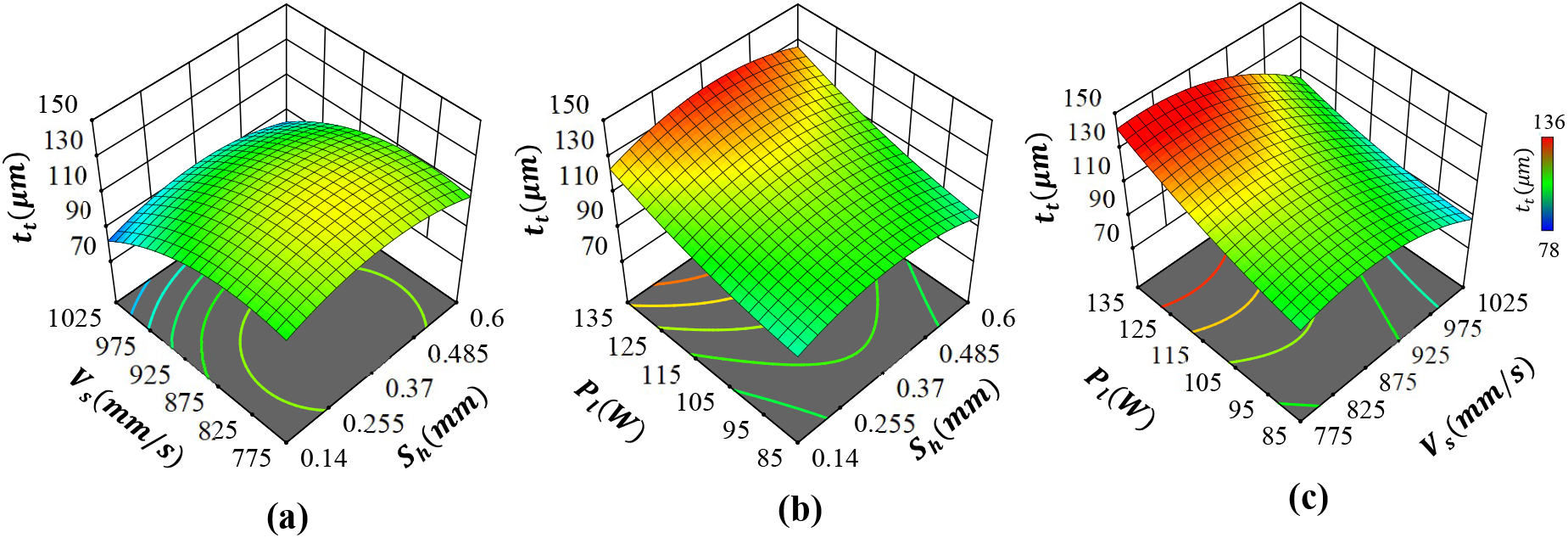
Influence of LPBF process parameters on Co-Cr-Mo track thickness showing interaction effects between (a) hatch spacing and scanning rate, (b) hatch spacing and laser power and (c) scanning rate and laser power.

As shown in Fig. 9c, the Co-Cr-Mo track thickness is primarily dictated by *P*_*l*_ and *V*_*s*_ with significant interaction effects between the two parameters. Therefore, achieving the thinnest track required careful control of both scanning rate and laser power. The strong interaction effects mean that the thinnest track was formed when both the laser power and scanning rate were at their lowest. As such, controlling either *P*_*l*_ or *V*_*s*_ without considering their interaction is unlikely to achieve the thinnest possible LPBF process-induced Co-Cr-Mo track. Deriving the order of influence, the most significant terms on *t*_*t*_ are the interaction effects of *V*_*s*_ and *P*_*l*_ in the order *P*_*l*_*V*_*s*_ > *V*_*s*_ > *P*_*l*_ with the least influence from *S*_*h*_. Consequently, to generate the finest porous Co-Cr-Mo architecture, a higher scan speed and lower laser power that induces sufficient energy density for the thinnest but fully melted track thickness is required.

#### 3.4.2. Pore diameter

The pore diameter is primarily dictated by the hatch spacing as evident from Fig. 10a with no influence from scanning rate or laser power as shown in Fig. 10b and Fig. 10c, respectively. The dependency of pore diameter on the hatch distance is also linear with the smallest and largest *ϕ*_*d*_ consistent with hatch spacing as shown in Fig. 10a and 10b. This response was expected as the parameter driving the distance between two adjacent tracks is hatch spacing. As shown in Fig. 10a and 10b, variations in *V*_*s*_ or *P*_*l*_ cannot introduce any changes to pore diameter signified by identical response surface. It can be also seen that there are no interaction effects between *V*_*s*_ and *P*_*l*_ as well demonstrated by the flat cure in Fig. 10c. As such the most significant term dictating the pore diameter of Co-Cr-Mo is the first-order effects of hatch spacing.

**Fig. 10.**
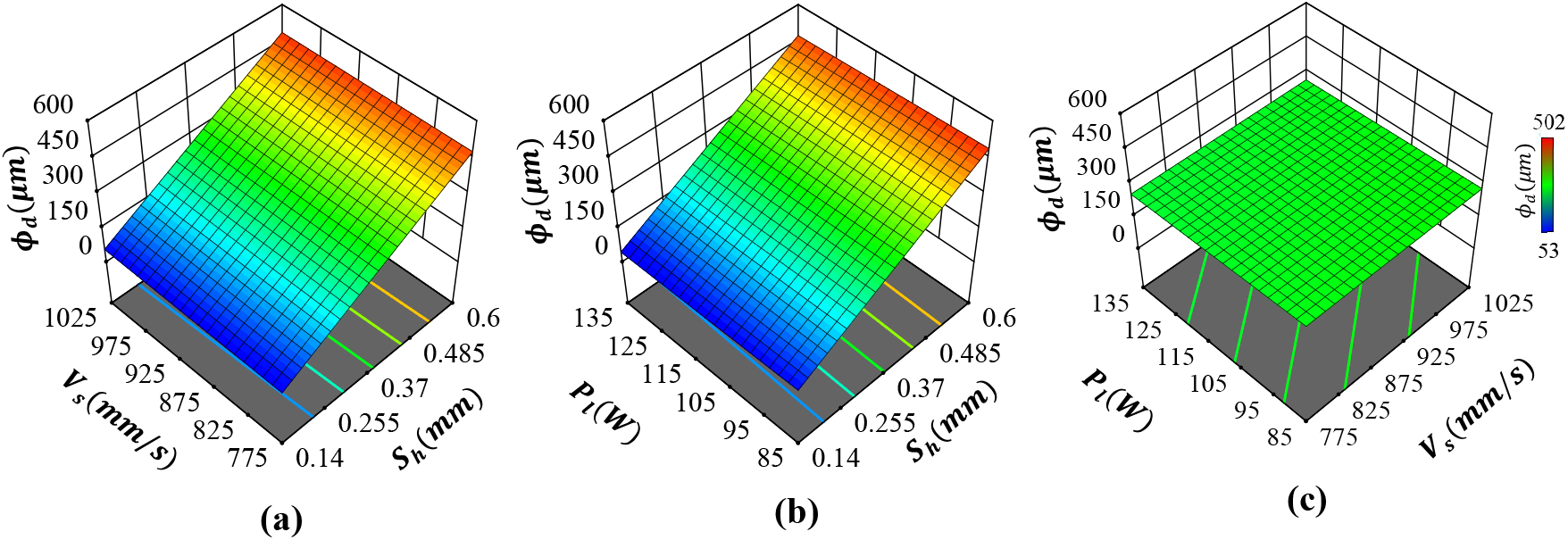
Influence of process parameters on the pore diameter of Co-Cr-Mo showing the interaction effects between(a) hatch spacing and scanning rate, (b) hatch spacing and laser power and (c) scanning rate and laser power.

The influence of process parameters on both track thickness and pore diameter reveals that the interaction effects can be used to simplify the Co-Cr-Mo microporous printing process. Overall, the targeted porosity can be achieved by carefully modulating the process parameters, which allows ease of customisability, and scalability. The influence of the process parameters on the material has also been analytically represented, which allows identifying the most optimal parametric combination based on the requirement. For the study under consideration a fine (<*t*_*t*_ and < *ϕ*_*d*_) architecture suitable for antiviral evaluation is the aim. The analysis so far shows that this can be achieved by minimising *P*_*l*_ and *S*_*h*_ while maximising *V*_*s*_ which results in thinner tracks as close as possible but not so close as to introduce a dense part.

### 3.5. Multi-objective optimisation

#### 3.5.1. Predicting optimal solution

The analysis so far has demonstrated the influence of the process parameters in the development of Co-Cr-Mo porous material. However, an accurate parametric combination that will lead to the best possible porous architecture is yet to be established. Doing this requires a multi-objective numerical description of the optimisation problem. To maximise the antiviral load, the optimum Co-Cr-Mo material should be porous in a way that maximises the surface area while featuring stable and consistent tracks. In other words, the optimised structure should allow for the smallest *ϕ*_*d*_ without producing a dense structure while featuring the thinnest *t*_*t*_. This means that the multi-objectives algorithm should look for a desirability criterion to minimise both *ϕ*_*d*_ and *t*_*t*_ to a non-zero positive integer. In this regard, the optimisation problem can be formulated as shown in Eq. (5):

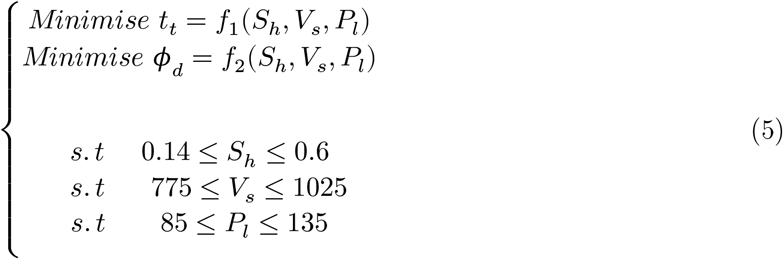

A multi-objective optimisation problem usually yields multiple solutions. As such a desirability function is used to combine them into a single objective (D) that meets the desirable range for each response (*d*_*i*_). The desirability approach is one of the most widely used methods in parametric optimisation where multiple responses are involved. It is based on the idea that the targeted outcome has multiple desired characteristics, any solution where the outcomes fall outside the desired limits is inadequate. Using this technique, the least and most desirable outcomes can be represented between 0 and 1 respectively. When *n* is the number of responses, the multi-objective function is a geometric mean of all transformed responses as shown in Eq. (6):

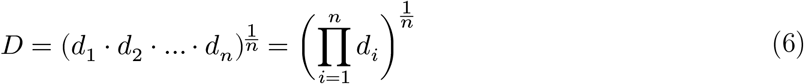

Fig. 11 shows the results of the multi-objective optimisation as a desirability function considering all interaction effects informed by the surrogate model. The highest desirability of 0.9 can be achieved at a scanning rate of 975-1025 mm/s, laser power of 85-105 W, and a hatch spacing of 0.14 mm. Before identifying the exact numerical values, it was important to consider particle size, which in this case ranged from 3-18 µm. This was set to prevent the powder particles from getting stuck in the pores leading to a dense part. As can be seen from Fig. 11a, despite keeping the other parameters constant increasing the laser power decreases the desirability. Overall, the lowest desirable solution is at the lowest scan speed, hatch distance, and highest laser power as shown in Fig. 11c. Table 6 shows one of the parametric combinations that offer the highest desirability that was used to fabricate the Co-Cr-Mo validation sample.

**Table 6.** Optimal LPBF parametric combination selected for Co-Cr-Mo validation build.

| Number | $S_h$ (mm) | $V_s$ (mm/s) | $P_l$ (W) | Desirability |
| --- | --- | --- | --- | --- |
| 1 | 0.14 | 1025 | 88 | 0.98 |

**Fig. 11.**
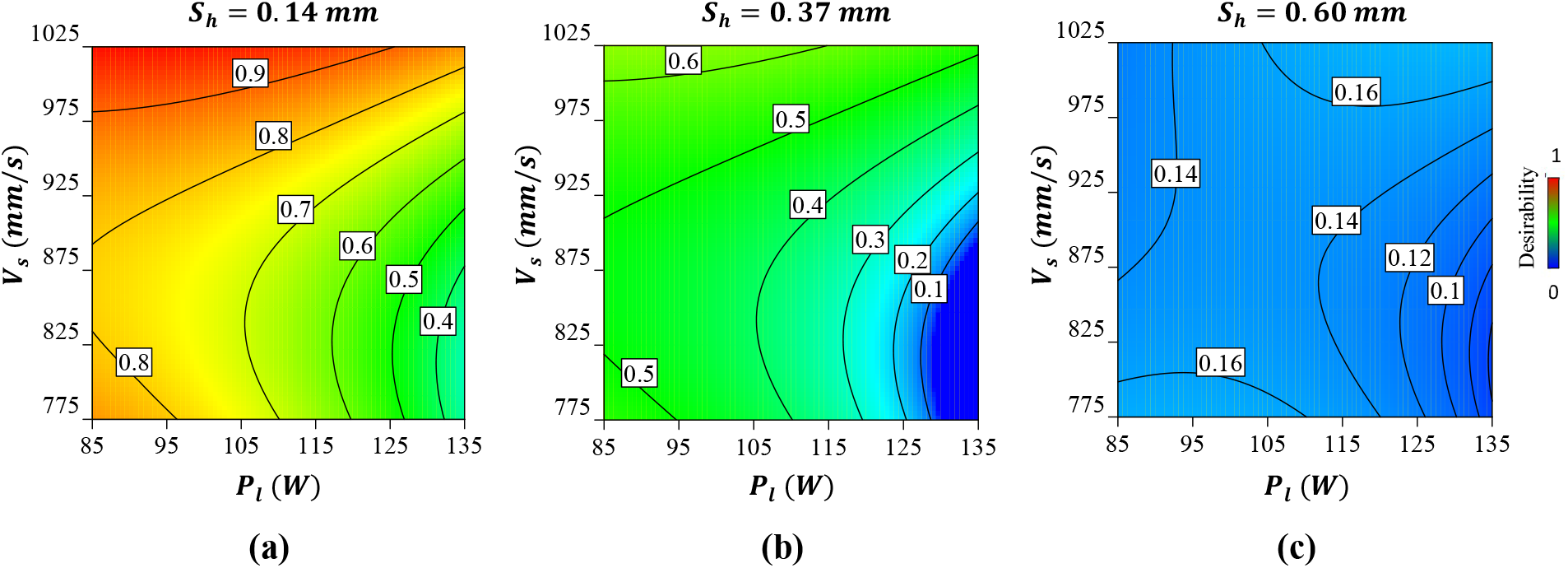
The desirability of the optimum solution for the Co-Cr-Mo microporous LPBF material showing (a) the effect of scanning rate and laser power at a hatch spacing of 0.14 mm (b) the effect of scanning rate and laser power at a hatch spacing of 0.37 mm and (c) effect of scanning rate and laser power at a hatch spacing of 0.60 mm. A desirability contour of 0 and 1 refer to the least and most optimum solution, respectively.

#### 3.5.2. Fabrication and validation of the optimised Co-Cr-Mo architecture

Co-Cr-Mo samples were additively manufactured using the predicted LPBF process parameter combinations listed in Table 6. The printed sample was analysed under SEM which confirms a fine porous architecture as shown in Fig. 12a. The magnified Fig. 12b confirms an even and consistent track and pore formation. Any further reduction in pore diameter runs the risk of powder particles becoming embedded in the porosity. As such the predicted optimum architecture by the surrogate model is closer to the potential porosity limit that can be achieved using the current particle and laser spot size.

**Fig. 12.**
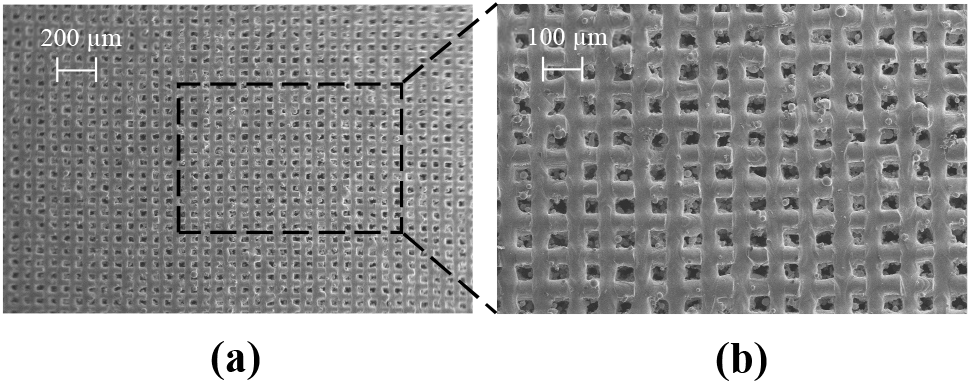
SEM data of as-built LPBF processed optimum Co-Cr-Mo microporous architecture showing (a) the overall pore distribution at a scale of 200 µm (b) a highlighted section showing the quality of the tracks and pores.

The results from the experimentally measured optimum Cu-W-Ag microporous material when compared to the surrogate model can be seen to be in good agreement as listed in Table 7. Overall, the surrogate model overestimated the track thickness and pore diameter by 3.9 % and % respectively. This small change between the predicted and experimented data can be attributed to the influence of particle size and partially melted powder at the track boundaries. Overall, the results show that the Co-Cr-Mo porous architecture with a feature size of 75 µm and pore diameter of 61 µm is achievable using LPBF, which transcends the current state-of-the-art feature size of 300 µm. As such, any further refinement in achievable pore size requires finer feedstock and further adjustments in laser modulation. Subsequently, the as-built Co-Cr-Mo optimised porous material was used for antiviral testing against the SARS-CoV-2 viral model to evaluate its antiviral effectiveness.

**Table 7.**
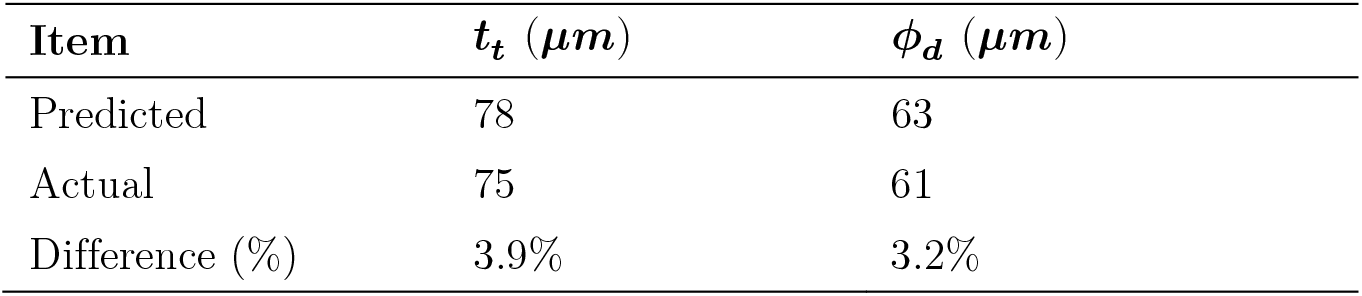
Comparison between actual and predicted values of the optimum Co-Cr-Mo microporous material.

| Item | $t_t$ ($\mu\text{m}$) | $\phi_d$ ($\mu\text{m}$) |
| --- | --- | --- |
| Predicted | 78 | 63 |
| Actual | 75 | 61 |
| Difference (%) | 3.9% | 3.2% |

### 3.6. Antiviral characterisation

The optimised microporous Co-Cr-Mo material was printed, and antiviral tests were performed (*n=3*) against the biosafe vrial model of SARS-CoV-2. The antiviral performance is summarised in Table 8, where the data at 30 minutes and 5 hours compared against a control sample, commercial porous fabric, and current state-of-the-art copper architecture. The corresponding visuals of the samples at 30 min. and 5 hrs. of contact with the phage are presented in Fig. 13. The control sample did not exhibit any antiviral activity at 30 mins. or 5 hrs. as shown in Fig. 13a and Fig. 13d, respectively. The commercial porous fabric, which was used as a reference material showed no antiviral performance at 30 mins. (Fig. 13b) or 5 hrs. (Fig. 13e) as expected. In comparison, the optimised Co-Cr-Mo LPBF porous architecture resulted in 100% viral inactivation at both contact times where no plaques can be observed as shown in Fig. 13c and 13f, respectively.

**Table 8.**
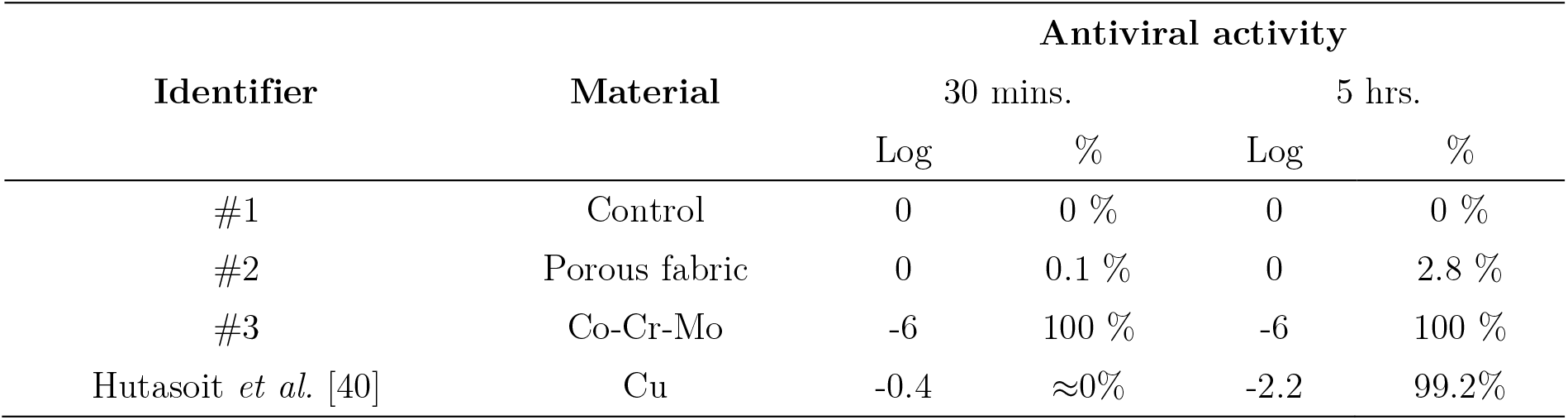
Antiviral performance against the viral model of SARS-CoV-2 of the control samples, Co-Cr-Mo porous material developed and state-of-the-art from literature.

**Fig. 13.**
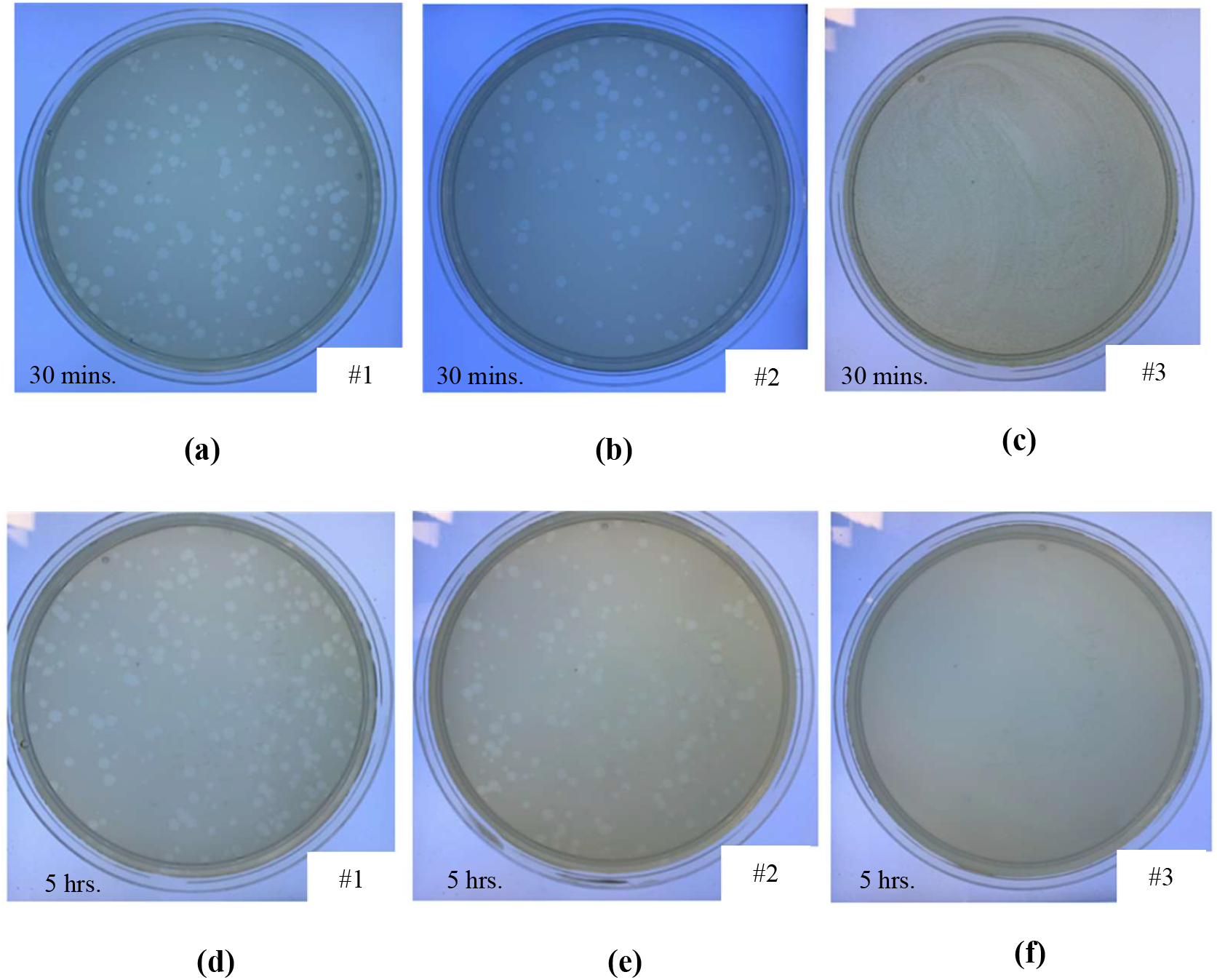
Loss of *phage phi 6* viability measured by the double-layer method at 10^−1^ dilution showing phage in contact with (a) control sample after 30 minutes, (b) commercial porous fabric after 30 minutes, (c) Co-Cr-Mo porous material after 30 minutes, (d) control sample after 5 hours (e) commercial porous fabric after 5 hours and (f) Co-Cr-Mo porous material after 5 hours.

The results revealed that the novel Co-Cr-Mo microporous architecture developed in this study have excellent antiviral properties (100% viral inactivation in 30 mins.) against phage phi 6 used as enveloped RNA viral model of SARS-CoV-2. The antiviral performance of the Co-Cr-Mo porous material is also superior to the current state-of-the-art copper [40], which showed 99.2% inactivation of SARS-CoV-2 in 5 hrs and no antiviral performance at 30 mins. as compared in Table 8. Generally, the SARS-CoV-2 virus can survive for 4-5 days in a range of materials [66] such as plastics, ceramics, stainless steel and glass as listed in Table 1. As such this paper presents the first metallic material that shows high (<30 mins.) antiviral performance against an enveloped virus like SARS-CoV-2 and influenza virus. This means that the material is suitable to be used as permanent antiviral devices aiding response against viral pathogens. Further development in this direction can aid pandemic preparedness to protect human beings in a more effective way while reducing the environmental impacts of disposable viral control devices in critical environments.

## 4. Conclusion

This study demonstrates the use of laser powder bed fusion for on-demand manufacturing of a novel Co-Cr-Mo porous material that shows superior antiviral activity outperforming common antiviral metals such as silver and copper. The paper reveals a surrogate model that allows porosity personalisation without the need for complex geometry data. This is achieved through controlling the LPBF process parameters such as laser power, scanning rate and hatch spacing. The proposed methodology simplifies the data requirement and pre-processing often required for printing porous materials. The surrogate model developed in this study showed that the most significant parameters for Co-Cr-Mo track thickness (*t*_*t*_) were the interaction effects of scanning rate (*V*_*s*_) and laser power (*P*_*l*_) in the order *P*_*l*_*V*_*s*_ > *V*_*s*_ > *P*_*l*_. For pore diameter (*ϕ*_*d*_), the hatch spacing (*S*_*h*_) has the most significant effect. The optimised Co-Cr-Mo microporous materials showed 100% viral inactivation against the phage phi 6 used as enveloped RNA viral model of SARS-CoV-2 in 30 minutes. The evolution of this and future pandemics will bring unexpected situations where the ability to print and personalise on-demand antiviral materials can achieve rapid solutions. Furthermore, the proposed methodology can be adopted to conceive functional antiviral materials that can be fabricated close to point-of-care.

## Acknowledgements

This research was conducted with support from the CALMERIC grant (European Commission, Grant number: 32R19P03053); University of Wolverhampton; Additive Analytics UK and EOS GmbH.

## Data availability

The data that supports the findings of this study are available from the corresponding author upon reasonable request.

